# Efficient APC/C substrate degradation in cells undergoing mitotic exit depends on K11 ubiquitin linkages

**DOI:** 10.1101/016139

**Authors:** Mingwei Min, Tycho Mevissen, Maria De Luca, David Komander, Catherine Lindon

**Affiliations:** University of Cambridge Department of Genetics, Cambridge, UK; Medical Research Council Laboratory of Molecular Biology, Cambridge, UK

## Abstract

The ubiquitin proteasome system (UPS) directs programmed destruction of key cellular regulators via post-translational modification of its targets with polyubiquitin chains. These commonly contain Lysine 48 (K48)-directed ubiquitin linkages, but chains containing atypical Lysine 11 (K11) linkages also target substrates to the proteasome, for example to regulate cell cycle progression. A single ubiquitin ligase, the Anaphase Promoting Complex/Cyclosome (APC/C), controls mitotic exit. In higher eukaryotes, the APC/C works with the E2 enzyme UBE2S to assemble K11 linkages in cells released from mitotic arrest, and these are proposed to constitute an improved proteolytic signal during exit from mitosis. We have tested this idea by correlating quantitative measures of *in vivo* K11-specific ubiquitination of individual substrates, including Aurora kinases, with their degradation kinetics tracked at the single cell level. We report that all anaphase substrates tested by this methodology are stabilized by depletion of K11 linkages via UBE2S knockdown, even if the same substrates are significantly modified with K48-linked polyubiquitin. Specific examination of substrates depending on the APC/C coactivator Cdh1 for their degradation revealed Cdh1-dependent enrichment of K11 chains on these substrates, while other ubiquitin linkages on the same substrates added during mitotic exit were Cdh1-independent. Therefore we show that K11 linkages provide the APC/C with a means to regulate the rate of substrate degradation in a co-activator-specified manner.

**Abbreviations:**

APC/C: Anaphase Promoting Complex/Cyclosome
DUB: deubiquitinase
SAC: spindle assembly checkpoint
UPS: ubiquitin-proteasome system
ZM: ZM447439

## Introduction

The specificity of substrate targeting by UPS enables precise remodelling of the protein landscape in many cellular processes. One such process is mitotic exit, initiated by destruction of mitotic cyclin (thereby decreasing CDK1 activity) and followed by a wave of degradation of other mitotic regulators, each with distinct timing and kinetics (Min and Lindon, 2012). All of these degradation events are mediated by the APC/C (Pines, 2011). A longstanding question in the field, then, is how this single ubiquitin ligase specifies temporal degradation for a large number of substrates. Part of the answer lies with the temporal specificity of APC/C’s coactivators, with Cdc20 active from prometaphase until its degradation during mitotic exit and Cdh1 from anaphase onwards (Pines, 2011). The coactivation mechanism is proposed to function partly via recruitment of substrates, and partly through enhancement of ubiquitination activity mediated by the APC/C (Barford, 2011), with recent studies attributing the latter role to enhanced E2 efficiency and stabilization of E2-APC/C interaction in the presence of coactivator (Brown *et al*., 2014; Chang *et al*., 2014; Kelly *et al*., 2014; Van Voorhis and Morgan, 2014).

In mammalian cells, APC/C is thought to act via the concerted activity of two different E2s. UBE2C adds the first ubiquitin or short ubiquitin chain to a substrate (‘priming’), while UBE2S attaches ubiquitin to the already attached ubiquitin molecules, elongating polyubiquitin chains in a K11 linkage-specific fashion (Williamson *et al*., 2009; Matsumoto *et al*., 2010; Wu *et al*., 2010; Wickliffe *et al*., 2011a). Linkage specificity of UBE2S accounts for an abrupt increase in K11 linkages seen in cells released from mitotic arrest, when the APC/C reaches its most active state (Matsumoto *et al*., 2010). It is therefore widely accepted that regulation of mitotic exit is one of the major roles of K11 chains (Bremm and Komander, 2011; Wickliffe *et al*., 2011b), even though UBE2S is not required for destruction of cyclin B1 in unchallenged mitotic exit (Garnett *et al*., 2009).

A recent study identified Nek2A as a Ube2S-dependent substrate of the APC/C in prometaphase (Meyer and Rape, 2014). Interestingly, the authors showed that Nek2A was modified with a variety of short chains by UBE2C, with UBE2S elongating short chains in a K11-linkage specific manner to generate branched chains. Such heterotypic chains were shown to constitute an improved degradation signal in prometaphase, when the APC/C is partially inhibited by the spindle assembly checkpoint (SAC) (Meyer and Rape, 2014). In mitotic exit however, once the SAC is silenced, there is a dramatic rise in K11-containing polyubiquitin conjugates. It is not known how these K11 linkages contribute to degradation of specific substrates as cells exit mitosis.

In this study we combine a cell-based ubiquitination assay that allows quantitative interrogation of ubiquitin conjugates on individual substrates (Min *et al*., 2013; Lee *et al*., 2014), with *in vivo* degradation assays that allow tracking of their degradation kinetics at the single cell level (Clute and Pines, 1999), to advance our understanding of the role and regulation of K11 linkages in mitotic exit.

## Results and discussion

### Abundance of K11 chains rises sharply in mitotic exit

The abrupt increase in K11 linkages in cells released from mitotic arrest suggests that K11 linkages could contribute to mitotic exit. However, whereas UBE2S is required for efficient mitotic progression following chronic drug-induced SAC activation, it is dispensable for mitotic exit in unchallenged mitosis (Garnett *et al*., 2009), raising the question of whether the boost of K11 linkages at mitotic exit also occurs during unperturbed mitosis. To test this, we synchronized U2OS cells at the G1/S phase boundary by double thymidine block and collected samples after release into fresh medium over a timecourse that followed progress through mitosis. The mitotic peak appeared 10 hours after release as indicated by phosphorylation of Histone H3 (Figure 1A). The decreasing level of Aurora A at 12 hours confirmed that mitotic exit had begun at this time, and was correlated with a sharp increase in the intensity of linkages detected using an antibody specific for K11-linked ubiquitin conjugates (Matsumoto *et al*., 2010) (Figure 1A). All material detected with this antibody in mitotic exit extracts was dependent on UBE2S (Figure 1B). We concluded that a dramatic increase in abundance of K11 chains accompanies mitotic exit, whether or not cells have previously undergone mitotic arrest.

**Figure 1.**
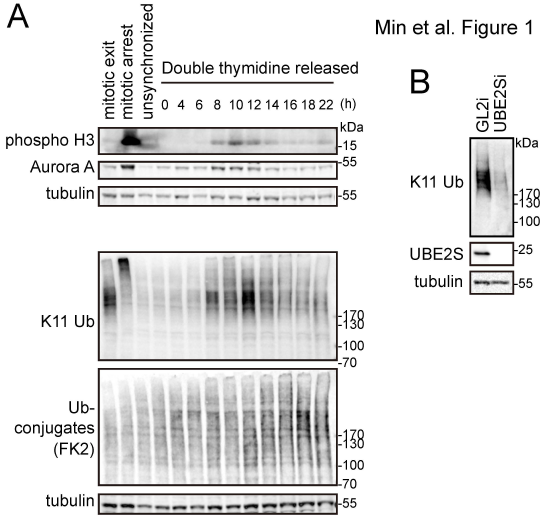
Abundance of K11 linkages rises strongly in unperturbed mitotic exit. **A** U2OS cells were synchronized at G1/S boundary using double thymidine block and then released into fresh medium. Samples taken at the indicated time points were blotted and probed with K11 linkage-specific antibody and other markers. For mitotic control samples, cells from double thymidine block were released into fresh medium containing 10 μ M STLC (S-trityl-L-cysteine) for 16 hr. SAC arrested cells were collected by mitotic shake-off, as mitotic arrest sample. AurB inhibitor ZM447439 was used at 10 μM to silence the SAC for 80 min to obtain the mitotic exit sample. **B** U2OS cells treated with siRNA against GL2 (control) or UBE2S were synchronized at mitotic exit and harvested 44 hr after transfection for immunoblotting with antibodies as indicated.

### APC/C substrates are modified with K11 ubiquitin linkages during mitotic exit

To investigate the function of K11 linkages during unperturbed mitotic exit, we sought to identify K11 acceptors by testing whether specific known anaphase substrates of APC/C-dependent proteolysis are modified with K11 linkages.

We have recently described a sensitive and robust cell-based assay for measuring ubiquitinated fractions of exogenously expressed GFP-tagged substrates in cells synchronized at mitotic exit, using drug-release protocols that generate a sharp boost in K11 linkages (Supplemental Figure S1). We applied this assay to two well-known anaphase substrates, mitotic kinases Aurora A and Aurora B (AurA and AurB). Following purification of AurA- and AurB-Venus from mitotic exit cells, we interrogated samples with linkage-specific K11 antibody and with GFP antibody. K11 linkages were clearly present on both Aurora kinases and abrogated by siRNA-mediated depletion of UBE2S (Ube2Si) (Figure 2A, B). Total ubiquitination on Aurora kinases was only partially affected, with approximately half of polyubiquitin conjugates remaining after Ube2Si (Figure 2A, C). Ubiquitin conjugates detected on these substrates were specific to mitotic exit (Supplemental Figure S2). We obtained quantifiably comparable results from stable, inducible cell lines (Floyd *et al*., 2013) or from cells transiently expressing AurA-Venus and AurB-Venus (Figure 2B, C). Quantifying the total ubiquitin signal as a function of molecular weight as shown in Figure 2D, we observed no apparent decrease in size of polyubiquitin conjugates after Ube2Si as would be expected if there were accumulation of shorter ubiquitin chain conjugates under these conditions. Therefore depletion of UBE2S did not appear to abolish the general elongation of ubiquitin chains.

**Figure 2.**
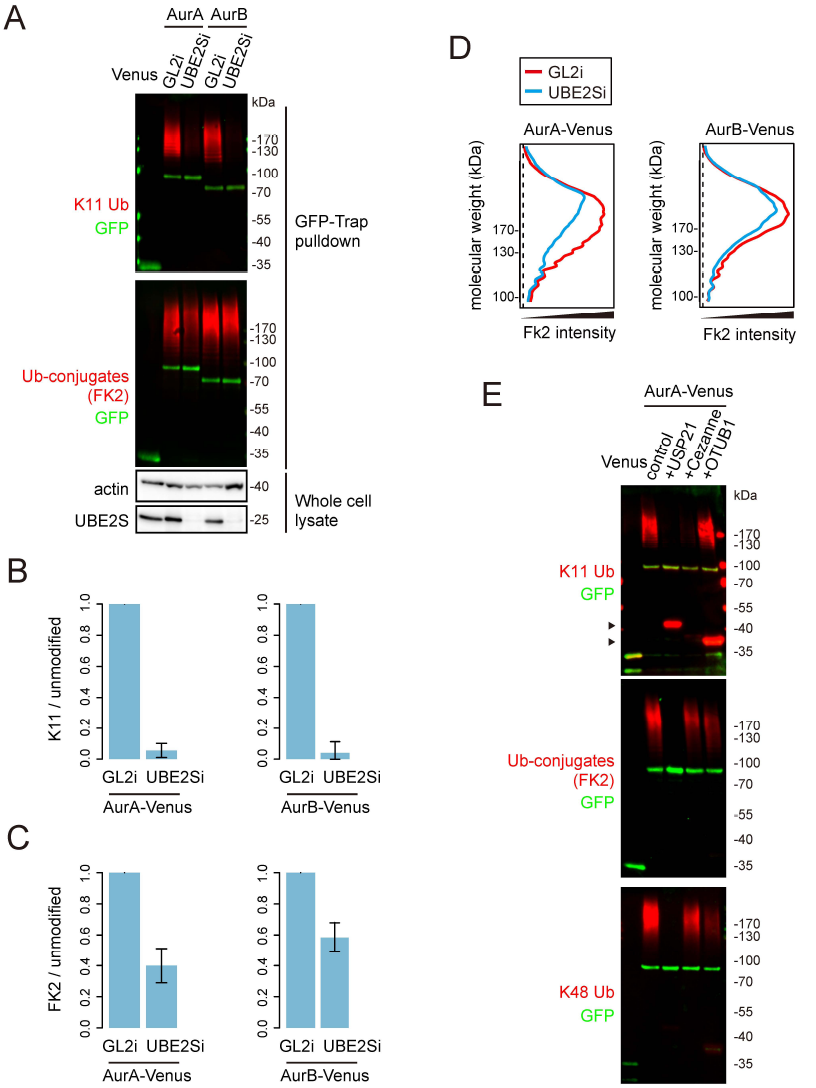
K11 chains are built on Aurora kinases at mitotic exit, in a UBE2S-dependent manner. **A-E** U2OS-AurA-Venus cells and U2OS-AurB-Venus cells were transfected with siRNA oligos against GL2 or UBE2S for 44 hr. Transfected cells were induced for expression of Venus-tagged substrates for 16 hr and synchronized to mitotic exit before harvesting for *in vivo* ubiquitination assays. **A** K11-specific antibody was used to signal K11 linkage and FK2 antibody for total ubiquitin linkages. GFP antibody recognises the Venus tag. **B-C** Quantifications of ubiquitin-modified to unmodified Aurora-Venus measured from **A** and a further two experiments, carried out with cells either inducibly or transiently expressing Aurora-Venus. In each case the ratio of modified:unmodified substrate was normalized to the corresponding GL2i sample. The bar chart shows the mean ± s.d. of the three independent experiments, showing ratios of K11-specific linkages (**B**) and total linkages (**C**). **D** Molecular weight distribution of ubiquitinated Aurora-Venus in control or UBE2S-depleted cells in the blots shown in **A**. **E** AurA-Venus purified as in **A** was subjected to DUB ‘restriction’ analysis. For this assay, AurA-Venus on beads was incubated with buffer alone, 1.5 μM USP21, 0.5 μM Cezanne, or 15 μM OTUB1 at 37 °C for 30 min as previously described (Mevissen *et al*., 2013). Arrowheads indicate unspecific staining of USP21 and OTUB1 by the K11 ubiquitin antibody.

We considered the possibility that there might exist compensatory mechanisms of ubiquitination to accommodate cells to UBE2S siRNA treatment, for example an increase in multi- versus poly-ubiquitin conjugates (Dimova *et al*., 2012), which would mask the loss of UBE2S-dependent K11 linkages. To test this idea, we used ubiquitin chain restriction (UbiCRest) analysis on AurA-Venus purified from untreated cells (Mevissen *et al*., 2013). Digesting with the non-linkage-specific DUB USP21 abolished all polyubiquitin conjugated to purified AurA-Venus (Figure 2E). Digesting with the K11-specific DUB Cezanne (Bremm *et al*., 2010) depleted the polyubiquitin fraction in a manner that resembled UBE2Si treatment, with all K11 linkages lost and total ubiquitin staining partially reduced across the entire molecular weight range (Figure 2E). Treatment with K48-specific DUB OTUB1 removed a significant fraction of linkages recognized by a K48-specific antibody, whilst having little effect on the amount of K11 linkage detected. For Nek2A-like ‘branched’ ubiquitination (Meyer and Rape, 2014), OTUB1 treatment should have decreased the quantity of K11 linkages detected. However, OTUB1 did not decrease K11 chains on AurA, suggesting that such chains are either of a distinct architecture, or exist as unbranched chains, although we cannot exclude that OTUB1 fails to hydrolyse K11-modified K48 chains efficiently. Regardless, knocking down UBE2S expression faithfully reflects specific removal of K11 linkages from AurA and is a valid route to interfere with K11 linkage formation in mitotic exit cells.

### K11 ubiquitin linkages promote substrate degradation in mitotic exit

Next, we investigated degradation of individual mitotic exit substrates under conditions of Ube2Si, using live cell imaging (Clute and Pines, 1999). We chose five substrates that are targeted by APC/C for proteolysis from anaphase onwards. These include substrates whose degradation exclusively depends on the coactivator Cdh1, namely AurA, AurB and Cdc6, as well as substrates that can be targeted by either Cdh1 or Cdc20, namely Plk1 and KIFC1 (Mailand and Diffley, 2005; Floyd *et al*., 2008; Clijsters *et al*., 2013; Min *et al*., 2014).

We expressed fluorescent protein-tagged substrates and traced their levels in single cells in asynchronously proliferating cultures. We then normalized fluorescence levels to anaphase onset and *in silico* synchronized cell traces. For each of the substrates analysed, the timing of degradation with respect to anaphase onset was not altered by depleting UBE2S. We observed decreased degradation rates for all substrates (Figure 3A, B), suggesting that K11 linkages are important in efficient degradation of mitotic exit substrates. Moreover, in the absence of UBE2S all substrates underwent degradation with similar kinetics despite distinct degradation kinetics under control conditions (Figure 3C, D), indicating that K11 chains specify accelerated rates of degradation in a substrate-dependent manner. This seems a key difference to budding yeast *S. cerevisiae*, where APC/C and its E2s Ubc1 and Ubc4 assemble K48 chains only (Rodrigo-Brenni and Morgan, 2007), and degradation of a variety of substrates was shown in a recent study to proceed at identical rates (Lu *et al*., 2014).

**Figure 3.**
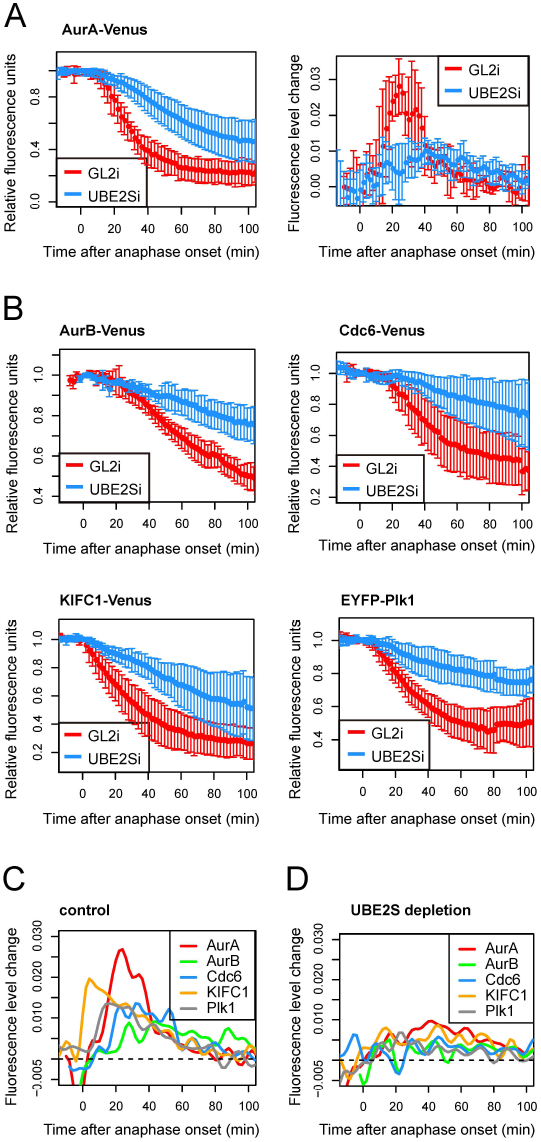
Efficient degradation of mitotic exit substrates requires UBE2S. **A-D** U2OS cells were transfected with indicated constructs together with control or UBE2S siRNA and imaged by fluorescence timelapse microscopy from 24 hr to 48 hr after transfection. Fluorescence intensity of Venus over time, in individual mitotic cells, was quantified and plotted as a function of anaphase onset. **A** *In vivo* degradation curve (lefthand panel) shows averaged intensities normalized to anaphase ± s.d. (n ≥ 5); degradation rate curve (righthand panel) shows the change in rate over time, derived from degradation curves as described in (Min *et al*., 2013). **B** Degradation curves for indicated substrates under conditions of GL2i or UBE2Si. **C** Degradation rate curves for mitotic exit substrates under control conditions. **D** Degradation rate curves for mitotic exit substrates after UBE2Si.

From an evolutionary viewpoint, K11 chains in higher eukaryotes may serve to specialize targeting of substrates to allow for more complex regulation of their proteolysis. For example only one Aurora kinase is present in yeast while there are three in mammals of which at least two are degraded during mitotic exit. Degradation of AurA starts soon after anaphase onset and both its degradation and activity are required for assembly of a robust spindle midzone (Floyd *et al*., 2008; Lioutas and Vernos, 2013; Reboutier *et al*., 2013). Beyond this point in mitotic exit, AurB activity is retained at the midbody to surveil abscission (Steigemann *et al*.) but otherwise needs to be degraded to promote respreading of the cell after mitosis (Floyd *et al*., 2013). These distinct roles in mitotic exit require that AurA and AurB disappear from the cell at different times. K11 linkages could therefore contribute to the temporal organization of substrate degradation through specifying different degradation kinetics.

### Regulation of K11 chain formation by APC/C-Cdh1

Given that the subset of mitotic exit substrates of cellular proteolysis could be major contributors to the K11-linked polyubiquitin detected in Figure 1A, we hypothesized that Cdh1 activity accounted for the dramatic increase in K11 chain assembly at mitotic exit. Indeed, we observed a 60% reduction in K11 linkage after siRNA-mediated Cdh1 depletion (Cdh1i) (Figure 4A), supporting the idea that K11 linkages could be the critical output of Cdh1 specificity in substrate degradation. Therefore we interrogated ubiquitin conjugates on individual substrates after Cdh1i. We found that, on the exclusively Cdh1-dependent substrates AurA and AurB (Floyd *et al*., 2008) (Supplemental Figure S3 A-D) Cdh1i treatment abolished K11 chains as effectively as Ube2Si (Figure 4B-C). By contrast, however, K11 chains on non-Cdh1-dependent substrate KIFC1 (Min *et al*., 2014) (Supplemental Figure S3 E-F), were not largely affected by Cdh1i (Figure 4D).

**Figure 4.**
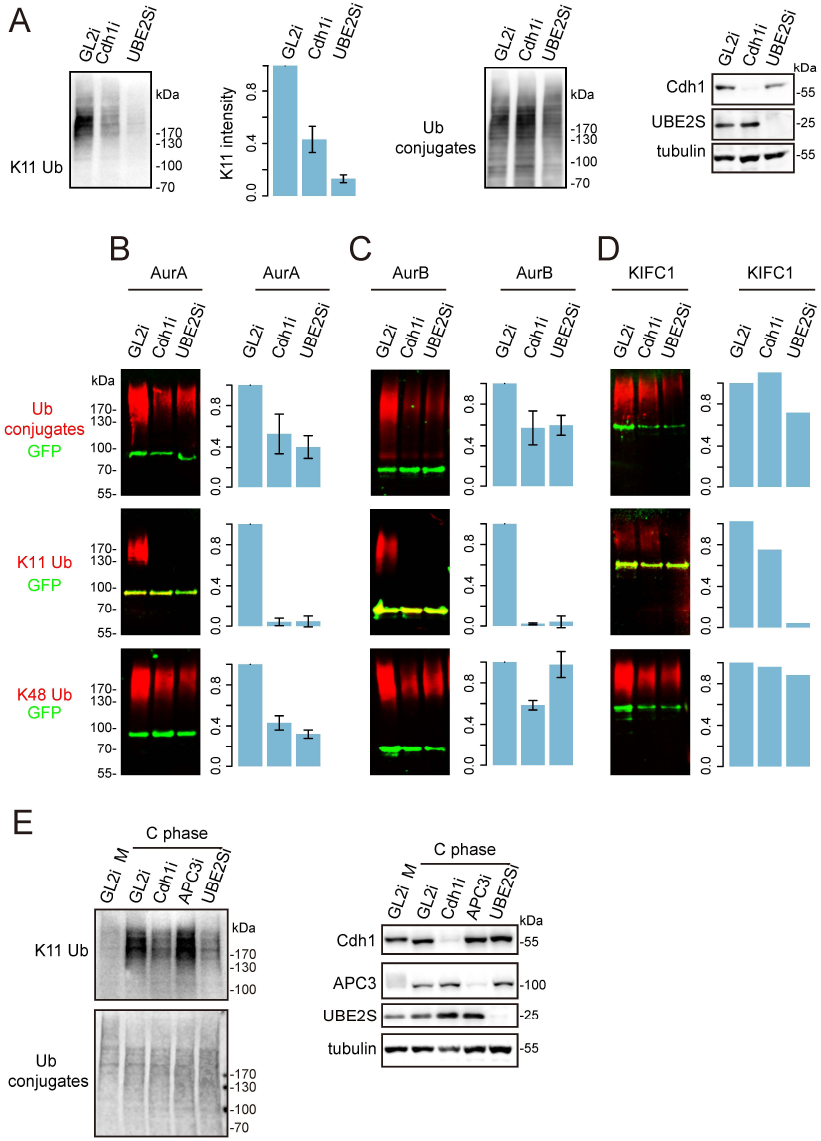
Regulation of K11 chain assembly by APC/C-Cdh1. **A** U2OS-^bio^Ub cells transfected with siRNA sequence against GL2 (control), Cdh1, or UBE2S were synchronized to mitotic exit. Whole cell lysate was interrogated by K11 linkage-specific antibody and by biotin antibody to show total ubiquitin conjugates. **B-D** U2OS-^bio^Ub cells were transfected with indicated Venus-tagged constructs together with control (GL2) or Cdh1 or UBE2S siRNA sequence, induced for ^bio^Ub expression for 44 hr and synchronized to mitotic exit. Cellular ubiquitination assays were carried out to detect total ubiquitin, K11 linkage and K48 linkage on each substrate. Bar plot shows mean measurements from three repeats for AurA and AurB and two repeats for KIFC1, with s.d. plotted where 3 repeats are available. *In vivo* degradation assays were carried out in parallel (shown in Supplemental Figure S3, together with representative blots to validate depletions. **E** U2OS-AurA-Venus cells were transfected with indicated siRNA and synchronized to prometaphase using a sequential thymidine and STLC block. Cells were then released into mitotic exit by treating with 300 nM CDK I/II inhibitor for 45 min before harvesting. Whole cell lysate were probed with indicated antibodies.

The coactivation mechanism is proposed to function partly via recruitment of substrates to the APC/C via specific motifs (Barford, 2011), and partly through enhancement of ubiquitination activity mediated by the APC/C, with recent studies attributing the latter role to enhanced E2 efficiency and stabilization of E2-APC/C interactions in the presence of coactivator (Brown *et al*., 2014; Chang *et al*., 2014; Kelly *et al*., 2014; Van Voorhis and Morgan, 2014). Our results confirm that substrate recruitment does not explain the role for Cdh1 in Aurora kinase degradation, since total polyubiquitin associated with Aurora kinases was only reduced by approximately 50%, with K48 chains still assembled on these substrates, in the absence of Cdh1 (Figure 4B-C). Therefore our results indicate that assembly of K11 chains, but not other linkages, correlates with substrate specificity in degradation conferred by Cdh1.

A recent study of coactivator function showed that Cdc20 plays a role in UBE2S activation by stabilizing the interaction between APC/C and UBE2S (Kelly *et al*., 2014). Cdh1 also binds UBE2S (Williamson *et al*., 2009) and our findings suggest that Cdh1 may act in a similar way. In the Kelly *et al*. study it was reported that mitotic exit APC/C was less active in stimulating UBE2S activity than APC/C^Cdc20^, based on measurement of UBE2S activity in the presence of Cdh1 and APC/C purified from synchronized cell extracts using APC3 antibody. APC3 is a core APC/C subunit required for cyclin B degradation in metaphase and proposed to interact with Cdh1 (Izawa and Pines, 2011; Chang *et al*., 2014). Strikingly, we found that depleting APC3 does not affect assembly of K11 linkage in mitotic exit, either in the whole cell lysate or on APC/C substrate AurA (Figure 4E). Therefore APC3 is not required for building K11 chains in mitotic exit. Although unexpected, this result can reconcile the finding from Kelly *et al*. that APC3-bound APC/C from mitotic exit cells is less active in building K11 linkages than that purified from mitotic arrested cells, with the observation that K11 linkages are several-fold more abundant in mitotic exit cells.

How would the process of stimulating K11 chain assembly couple with the substrate selectivity of Cdh1? K48 chains can still be assembled on Aurora kinases upon Cdh1 depletion, implying that these substrates are able to receive K48-linked ubiquitin chains from Cdc20-activated APC/C. Indeed, Cdc20 and Cdh1 can both bind to the canonical D-box and KEN box motifs (Barford, 2011). It has recently been shown that binding of coactivator increases the affinity of APC/C for an E2 (Chang *et al*., 2014) and, importantly, that in the case of the *S. cerevisiae* elongating E2 Ubc4, Cdh1-bound substrate further enhances E2 activity (Van Voorhis and Morgan, 2014). Therefore, we propose that substrate-specific stimulation promotes coactivator-dependent recruitment of UBE2S. The study from Barford and colleagues lends support to this idea by showing that Cdh1 binding induces displacement and flexibility of the APC/C catalytic module APC2-APC11 (Chang *et al*., 2014). Co-activator specificity in substrate degradation, mediated by K11 linkages, would by this account arise from coactivator-dependent positioning of the K11-specific machinery at the right proximity to substrates.

In conclusion, we show that the amount of K11 ubiquitin linkage in the cell is strongly cell cycle-dependent, with its peak at mitotic exit. K11 chains are most likely built on the majority of mitotic exit substrates. The distinct kinetics that we observe for different mitotic exit substrates are all modulated by K11 linkages, which are essential for the rapid degradation of some substrates, such as Aurora A and KIFC1. Moreover, K11 chain formation seems to correlate with substrate specificity conferred by Cdh1, providing a potential link between co-activator specificity in targeting and efficient substrate degradation in mitotic exit. We also demonstrate the importance of investigating ubiquitination in a cellular context. Although *in vitro* studies have generated extensive knowledge of the versatile enzyme activities in the UPS, tracking their activity in cells can reveal unexpected detail in how they contribute to cellular functions.

## Materials and Methods

### Cell culture and synchronization

U2OS-^bio^Ub cells and U2OS parental cells were described before in (Min *et al*., 2014) and U2OS-AurB-Venus in (Floyd *et al*., 2013). A tetracycline-regulated U2OS-AurA-Venus cell line was created from the same parental line as the U2OS-AurB-Venus cells, using pTRE-AurA-Venus plasmid and established procedures (Floyd *et al*., 2013). All U2OS cells were cultured in high glucose DMEM (GE healthcare). Cell culture medium was supplemented with FBS (10%), Penicillin-Streptomycin, amphotericin B, 500 μg/mL geneticin (all from PAA Laboratories) and 1 μg/mL tetracycline hydrochloride (Calbiochem). Tetracycline was removed from medium to induce expression of the corresponding construct. Synchronization at mitotic exit was achieved using a sequential thymidine block (20 hr, 2.5 mM) and STLC block (16 hr, 10 μM after 3 hours release from thymidine). Prometaphase cells were collected by mitotic shake-off and forced into mitotic exit by silencing the SAC using AurB inhibitor ZM447439 (10 μM) for 70 min, unless specified otherwise. Double thymidine synchronization experiment, cells were first treated with 2.5 mM thymidine for 20 hr, released into fresh medium for 12 hr. Then 2.5 mM thymidine was added into the culture for 12 hr before releasing again.

### Plasmids, siRNA and transfection

pVenus-AurB (Floyd *et al*., 2013), pEYFP-Plk1 (Lindon and Pines, 2004), pVenus-AurA and pVenus-KIFC1 (Min *et al*., 2014) were as described before. Cloning details for pTRE-AurA-Venus are available on request. pVenus-Cdc6 was a kind gift from Rob Wolthuis (Clijsters *et al*., 2013). siRNA sequence targeting APC3, Cdh1 and UBE2S were as previously described (Floyd *et al*., 2008; Garnett *et al*., 2009; Izawa and Pines, 2011). Plasmids and siRNAs were electroporated into cells using the Neon transfection system (Life Technology) using a program with voltage of 1150V, width of 30 ms and 2 pulses. Each transfection was carried out with 2X10^6^ to 10^7^ cells with 5 μg plasmid per 100μL transfection or 1 μM siRNA oligo in the transfection suspension, respectively.

### Cellular ubiquitination assay

To prepare one GFP-Trap pulldown sample for immunoblotting, about 5X10^6^ cells expressing a Venus-tagged substrate were lysed in 100 μL lysis buffer (50 mM Tris– HCl pH 7.6, 150 mM NaCl, 1 mM EDTA, 1% (v/v) triton; 1X protease inhibitor cocktail (Roche); 1 mM NaF; 1 mM Na_3_VO_4_; 50 mM *N*-ethylmaleimide). Following dilution with 900 μL dilution buffer (10 mM Tris–HCl pH 7.6, 150 mM NaCl, 0.5 mM EDTA, 1X protease inhibitor cocktail; 1 mM NaF, 1 mM Na_3_VO_4_, 50 mM *N*-ethylmaleimide), the lysate was cleared by centrifugation, applied to 5 μL pre-washed GFP-Trap_A beads and incubated for 2 hr at room temperature. The beads were then washed with 8 M Urea, 1% (w/v) SDS for 1 min, 10 mM Tris-HCl pH 7.6, 1M NaCl, 0.5 mM EDTA, 1% (v/v) triton for 3X5 min on roller and briefly rinsed in 1% (w/v) SDS. The beads were then boiled in 5 μL 4X sample buffer (Life Technology) containing 100 mM DTT for 10 min at 95 °C and the eluted sample was separated from the beads by centrifugation at 14000 g. Blots were probed with GFP antibody (with a IRDye fluorescent secondary antibody) and indicated ubiquitin antibodies (with a HRP-linked secondary antibody) for quantitative detection of unmodified and ubiquitinated Venus-tagged substrate, respectively, using a Li-Cor Odyssey^®^ Fc. Ubiquitination levels are indicated by the ratio of ubiquitin antibody intensity to GFP antibody intensity.

### Ubiquitin smear distribution profiling

The indicated ubiquitin blots were exported from Li-Cor Image studio as a non-saturated 8 bit image. For each lane, 3 line scans (plot profiles) were measured in Image J. The averaged profiles were generated in R using an in house script.

### In vivo degradation assay

Imaging and analysis were performed as described before (Min *et al*., 2013; Min *et al*., 2014).

### Antibodies

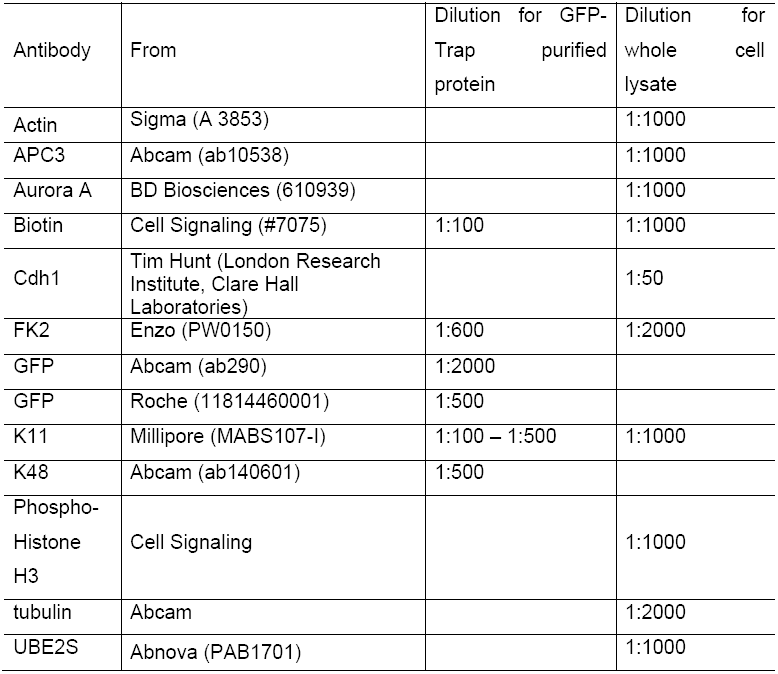

| Antibody | From | Dilution for GFP-Trap purified protein | Dilution for whole cell lysate |
| --- | --- | --- | --- |
| Actin | Sigma (A 3853) |  | 1:1000 |
| APC3 | Abcam (ab10538) |  | 1:1000 |
| Aurora A | BD Biosciences (610939) |  | 1:1000 |
| Biotin | Cell Signaling (#7075) | 1:100 | 1:1000 |
| Cdh1 | Tim Hunt (London Research Institute, Clare Hall Laboratories) |  | 1:50 |
| FK2 | Enzo (PW0150) | 1:600 | 1:2000 |
| GFP | Abcam (ab290) | 1:2000 |  |
| GFP | Roche (11814460001) | 1:500 |  |
| K11 | Millipore (MABS107-I) | 1:100 – 1:500 | 1:1000 |
| K48 | Abcam (ab140601) | 1:500 |  |
| Phospho-Histone H3 | Cell Signaling |  | 1:1000 |
| tubulin | Abcam |  | 1:2000 |
| UBE2S | Abnova (PAB1701) |  | 1:1000 |

## Acknowledgements

Our thanks to Chiara Marcozzi and Ben Kinnersley for help in setting up Aurora B degradation assays and to Rob Wolthuis for Cdc6-Venus. Yu Ye and Rhys Grant contributed valuable discussions during the course of this study. Work in CL lab was funded by Medical Research Council [G120/892], Cancer Research UK [C3/A10239] and the Department of Genetics. Work in DK lab is funded by Medical Research Council [U105192732], European Research Council [309756], and the Lister Institute for Preventive Medicine. MM was supported by Great Britain China Centre Educational Trust and the Henry Lester Trust. TM is funded by Marie Curie Initial Training Network “UPStream”.

## Author contributions

The study was conceived, designed and written by MM and CL. MDL generated the Aurora A-Venus cell line used in the study. Other experimental work was carried out by MM excepting UbiCRest assays carried out by TM. DK interpreted data and contributed to the manuscript.

---

**Figure S1.**
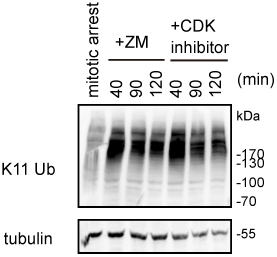
K11 linkages are enriched in mitotic exit under all conditions tested. U2OS cells were synchronized at prometaphase using a sequential thymidine (2.5 mM, 20 hr) and STLC (10 μM, 16 hr) block, collected by mitotic shake-off and released into mitotic exit using either ZM447439 at 10 μM or CDK I/II inhibitor at 300 nM for indicated time. Immunoblotting of cell extracts showed strong enrichment of K11 linkages as in Figure 1A.

**Figure S2.**
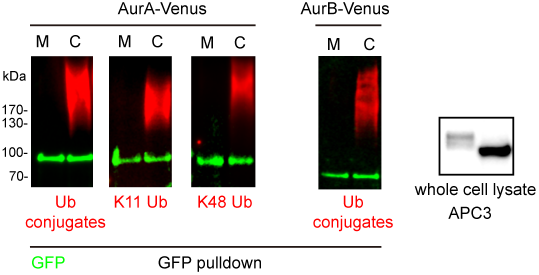
(related to Figure 2) Ubiquitin chains on AurA and AurB detected in our cellular ubiquitination assays were highly specific to mitotic exit. AurA-Venus cells and AurB-Venus cells were synchronized to prometaphase using a sequential thymidine and STLC block and collected by mitotic shake off. Half were harvested as the mitotic arrest (M) sample and the remainder released using ZM before harvesting, for the mitotic exit (C phase) sample. Immunoblot of cell lysates using antibody against APC3 shows the dephosphorylation-dependent increase in mobility of APC3 that provides a marker for mitotic exit. Venus-tagged Aurora kinases were purified for ubiquitination assays to show that all ubiquitination linkages detected on AurA- and AurB-Venus (in our assays) are specific to mitotic exit.

**Figure S3.**
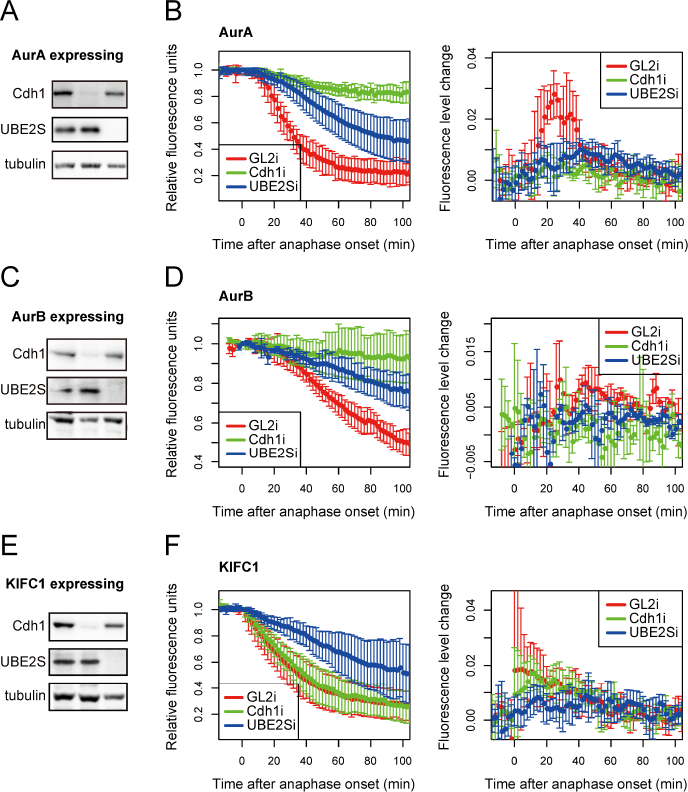
(related to Figures 3 and 4) Degradation of AurA and AurB are sensitive to Cdh1 depletion whilst degradation of KIFC1 is not. Cells co-transfected with Venus-tagged substrates and siRNA oligos against GL2 (control), Cdh1 or UBE2S, from the same experiments shown in Figure 4, were filmed by fluorescence timelapse microscopy for the substrate degradation assays described in Figure 3. **A, C, E**, validation of siRNA-mediated depletions by immunoblot; **B, D, F**, degradation curves and degradation rate curves for the indicated substrates. GL2i and UBE2Si curves are generated from the data used for Figure 3.

